# Amphotericin B Resistance in *Lomentospora prolificans* is associated with a soluble cell wall component

**DOI:** 10.64898/2026.06.25.734450

**Authors:** Nina T. Grossman, Arturo Casadevall

## Abstract

**Introduction:** *Lomentospora prolificans* is a pathogenic filamentous fungus that causes disease primarily in people with severely compromised immune systems. It is pan-resistant to antifungal drugs, but the mechanism of its resistance to amphotericin B (AMB) is unknown.

**Objectives:** We aimed to investigate the mechanism of resistance to AMB of *L. prolificans*.

**Methods:** The AMB susceptibility of *L. prolificans* protoplasts was measured using broth microdilution. *L. prolificans*, either intact, homogenized or fractionated was incubated with AMB in broth. The same activity was carried out with *Aspergillus fumigatus* as a control. This broth was then used to prepare microdilution plates with *Saccharomyces cerevisiae* to determine the activity of the conditioned AMB.

**Results:** AMB was 16-fold more effective in inhibiting the growth of *L. prolificans* protoplasts than conidia, but only two-fold more effective against *A. fumigatus* protoplasts than conidia. Incubation of *L. prolificans* hyphae with AMB in media diminished drug activity to a much greater extent than *A. fumigatus*, with 8-fold greater fungal mass of the latter required to achieve the effect of the former. Homogenization and fractionization of *L. prolificans* revealed that the factor inhibiting AMB activity was soluble with a mass >100 kda. DNase, trypsin, proteinase K, amyloglucosidase, SDS and 0.22 μm had no effect on the AMB resistance factor, while treatment with urea, acetonitrile inactivated it.

**Conclusion:** We report a different mechanism for AMB resistance based on the existence of a substance residing in the *L. prolificans* cell wall that can eliminate the antifungal activity of AMB.

## Introduction

*Lomentospora prolificans* is a pathogenic filamentous fungus that causes disease primarily in people with severely compromised immune systems. While *L. prolificans*-related disease has a fairly low incidence, with one study estimating 0.53 cases per million people annually in Australia (1), such cases have remarkably high fatality rates. Among patients with malignancy, which is a common risk factor for invasive lomentosporiosis, all-cause mortality can be as high as 85.7% (2). The primary reason for the deadly nature of *L. prolificans* infections is that the organism is pan-resistant to antifungal drugs, making it very difficult to treat (3, 4). *L. prolificans* represents one of the few fungal species that show inherent resistance to all three major classes of antifungal drugs – azoles, echinocandins and polyenes. An analysis of mostly clinical *L. prolificans* isolates found that the minimum concentration of drug necessary to successfully inhibit the growth of all but the most resistant 10% of the isolates was greater than the highest concentration of drug tested for all eight of the antifungals tested (4). In this characteristic, it surpasses even isolates of the notoriously resistant *Candida auris*, 90, 30 and 5% of which are resistant to fluconazole, amphotericin B (AMB) and echinocandins, respectively (5). However, only one mechanism of resistance has thus far been identified in *L. prolificans* – an FKS1 protein sequence that prevents efficient binding by echinocandin drugs (6, 7).

Among most human fungal pathogens, AMB resistance is rare and poorly understood. The mechanisms associated with reduced susceptibility to amphotericin B tend to fall into three broad categories. The first and most reported of these is alteration to the lipid composition of the cell membrane, generally from loss-of-function mutations in genes in the ergosterol synthesis pathway. This results in lower cell membrane ergosterol content, thereby providing less target for the drug to bind (8). The second category of AMB resistance mechanisms is the increased production of substances capable of binding amphotericin B and preventing it from reaching the cell membrane. Such substances include β-glucans, which is found in the extracellular matrix of *Candida* biofilms and appears to bind not just AMB, but also fluconazole (9–12), and melanin, which is hypothesized to sequester AMB away from the cell membrane and also impede the drug’s access to its target by reducing cell wall pore size in *Cryptococcus neoformans* (13–16). Finally, the third set of hypothesized mechanisms of AMB resistance compensates for the oxidative damage done by the drug, either by overexpression of catalase or upregulation of heat shock proteins (17–19).

As incomplete as the understanding of mechanisms of amphotericin B activity and resistance is, the current knowledge on mechanism of amphotericin B resistance in planktonic *L. prolificans* is even sparser. Two publications document successful attempts to interfere with the capacity of *L. prolificans* to melanize, the first by random mutagenesis and the second through the creation of knockouts of three genes in the melanin synthesis pathway (20, 21). Both groups found that the mutants remained resistant to amphotericin B treatment, even in the absence of melanin. Another study shows increases in SOD and catalase activity in response to AMB exposure in *L. prolificans*, as well as *Scedosporium minutisporum*, and *Scedosporium aurantiacum*, the latter of which shows much lower rates of resistance than *L. prolificans*, suggesting that SOD and catalase activity alone are probably not responsible for the remarkable resistance of *L. prolificans* (4, 22). Additionally, amphotericin B synergizes with echinocandins against *L. prolificans* (23, 24), suggesting that resistance to the former may be related to the cell wall, synthesis of which is inhibited by echinocandins.

In this paper, we sought to understand the potential role of the cell wall in the resistance of *L. prolificans* to amphotericin B. After noting that removal of the cell wall rendered *L. prolificans* protoplasts susceptible to AMB, we established that a soluble component of the *L. prolificans* cell wall could remove or inactivate AMB in solution.

## Results

To determine whether the cell wall contributes to antifungal resistance in *L. prolificans*, we examined the effect of cell wall removal on its susceptibility to AMB. Using a broth microdilution plate, we measured the minimum inhibitory concentration (MIC) of AMB against protoplasts and intact conidia of both *L. prolificans* and *A. fumigatus*. We found that AMB was 16-fold more effective against *L. prolificans* protoplasts than against its conidia (MIC 1 μg/ml vs 16 μg/ml, respectively, Table 1). In contrast, *A. fumigatus* showed only one dilution of difference between protoplasts and conidia (MIC 0.5 μg/ml vs 0.25 μg/ml). This suggested that the *L. prolificans* cell wall was essential to its unusual ability to grow in high concentrations of AMB, and that the capacity of the cell wall to augment AMB resistance was species-specific, rather than a common characteristic amongst filamentous fungi.

**Table 1.**
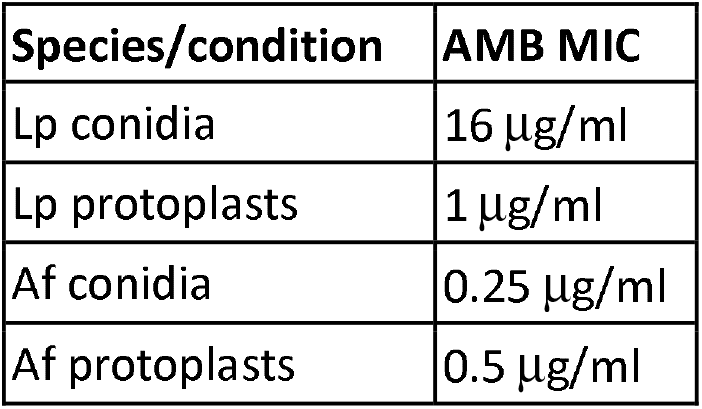
AMB susceptibility of *L. prolificans* (Lp) and *A. fumigatus* (Af) conidia and protoplasts by broth microdilution plate.

| Species/condition | AMB MIC |
| --- | --- |
| Lp conidia | 16 µg/ml |
| Lp protoplasts | 1 µg/ml |
| Af conidia | 0.25 µg/ml |
| Af protoplasts | 0.5 µg/ml |

We theorized that the *L. prolificans* cell wall could be acting as a drug sink – binding the AMB and preventing it from reaching the cell membrane. To test this hypothesis, *L. prolificans* hyphal mass was added to RPMI broth with AMB in various concentrations, shaken for an hour at 37°C, then filtered out. This sterile conditioned media was then used to prepare a broth microdilution drug susceptibility plate with a Saccharomyces cerevisiae strain of known AMB susceptibility (Figure 1). Accordingly, the MIC of the S. cerevisiae was used to determine how much drug activity remained in the media after incubation with *L. prolificans*, rather than the drug susceptibility of the yeast. Adding increasing concentrations of *L. prolificans* to the media diminished AMB activity in a dose-dependent manner, with 0.5, 1.0 and 1.5 mg/ml of fungal mass roughly multiplying the initial concentration of AMB necessary to inhibit yeast growth by on average two, four and eight, respectively (Figure 2). For comparison, the same process was conducted with *S. apiospermum* (MIC = 2 μg/ml), and *A. fumigatus*. While the addition of either of these fungi was sufficient to diminish the drug activity in the conditioned media, a much greater mass of both fungi was required to achieve the same effect as conditioning with *L. prolificans*. While 0.5 mg/ml of *L. prolificans* fungal mass reduced the drug activity by one dilution, 1.5 mg/ml of *S. apiospermum* and 4 mg/ml of *A. fumigatus* was necessary before the same reduction in drug activity was observed.

**Figure 1.**
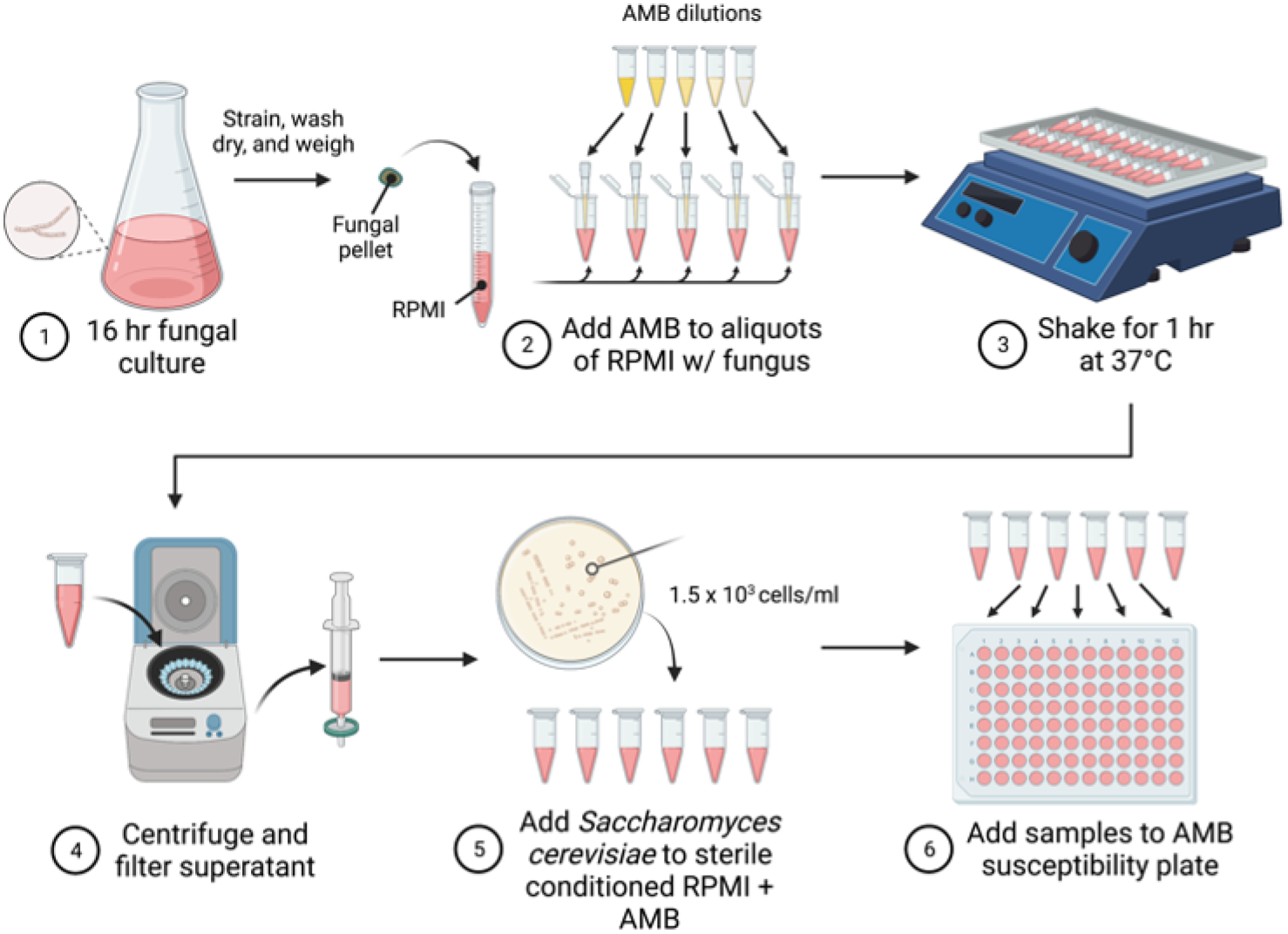
Steps involved in preparing the drug activity assay plate. Figure created using biorender.com.

**Figure 2.**
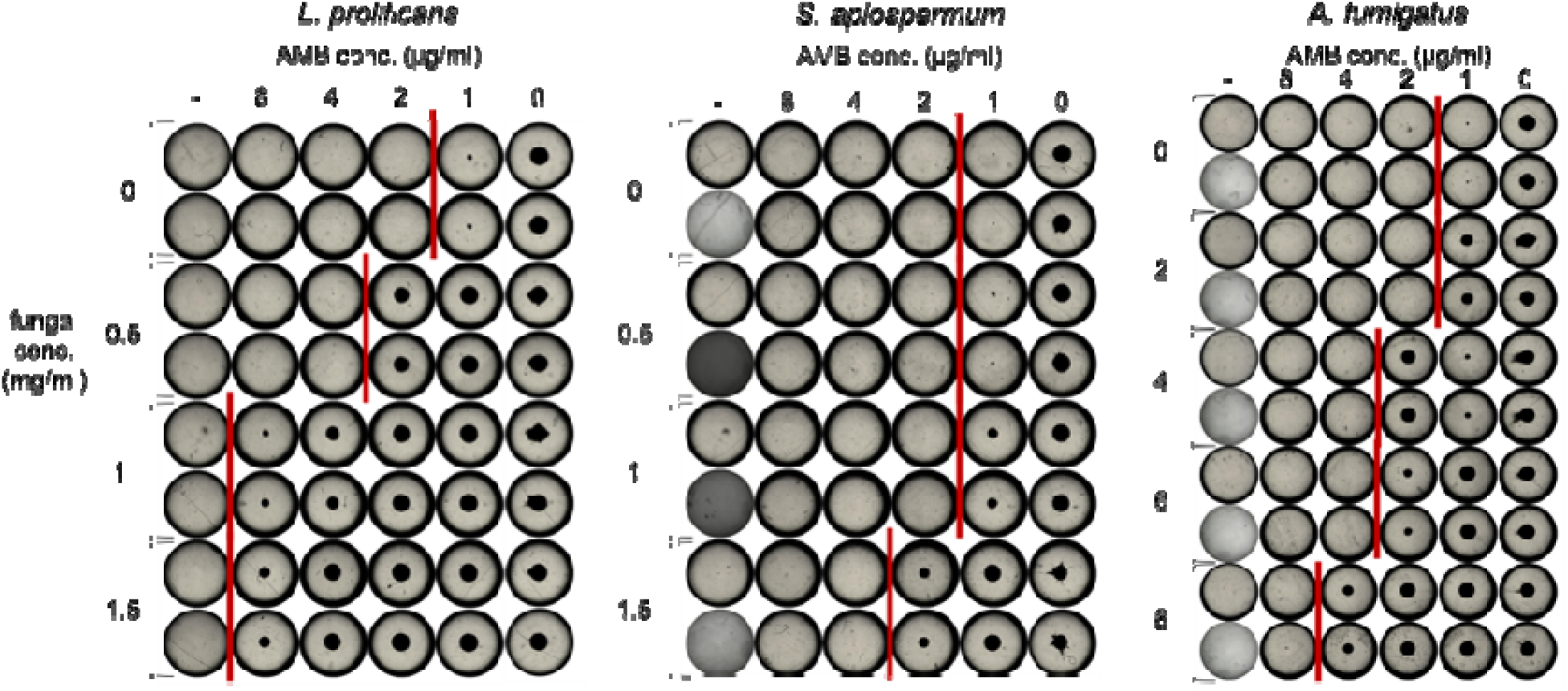
Reduction in AMB activity against Saccharomyces cerevisiae from incubation with intact *L. prolificans, S. apiospermum* and *A. fumigatus* hyphae. Various concentrations by weight of the listed fungi were shaken with AMB in RPMI before the media was sterilized and used in AMB susceptibility plate inoculated with S. cerevisiae. Growth beyond that shown in fungal-free controls (denoted by ‘-’) indicates depletion of drug activity. Red lines separate wells with growth from wells without growth. Similar results were obtained in five biological replicates in technical duplicates for *L. prolificans* and three biological replicates in technical duplicates for *S. apiospermum* and *A. fumigatus*.

Having established that the presence of *L. prolificans* reduced AMB activity to an extent not observed with other filamentous fungal pathogens that are more susceptible to AMB, we next explored whether this effect was contingent on direct contact between the fungus and the media. Accordingly, the drug activity assay was repeated with the alteration that the AMB was not added until after the fungus had been filtered out, such that the media was conditioned, but the drug was not. Under these conditions, no reduction of drug activity was observed, indicating that the drug was interacting with the fungus itself, rather than with a secreted factor (Table 2).

**Table 2.**
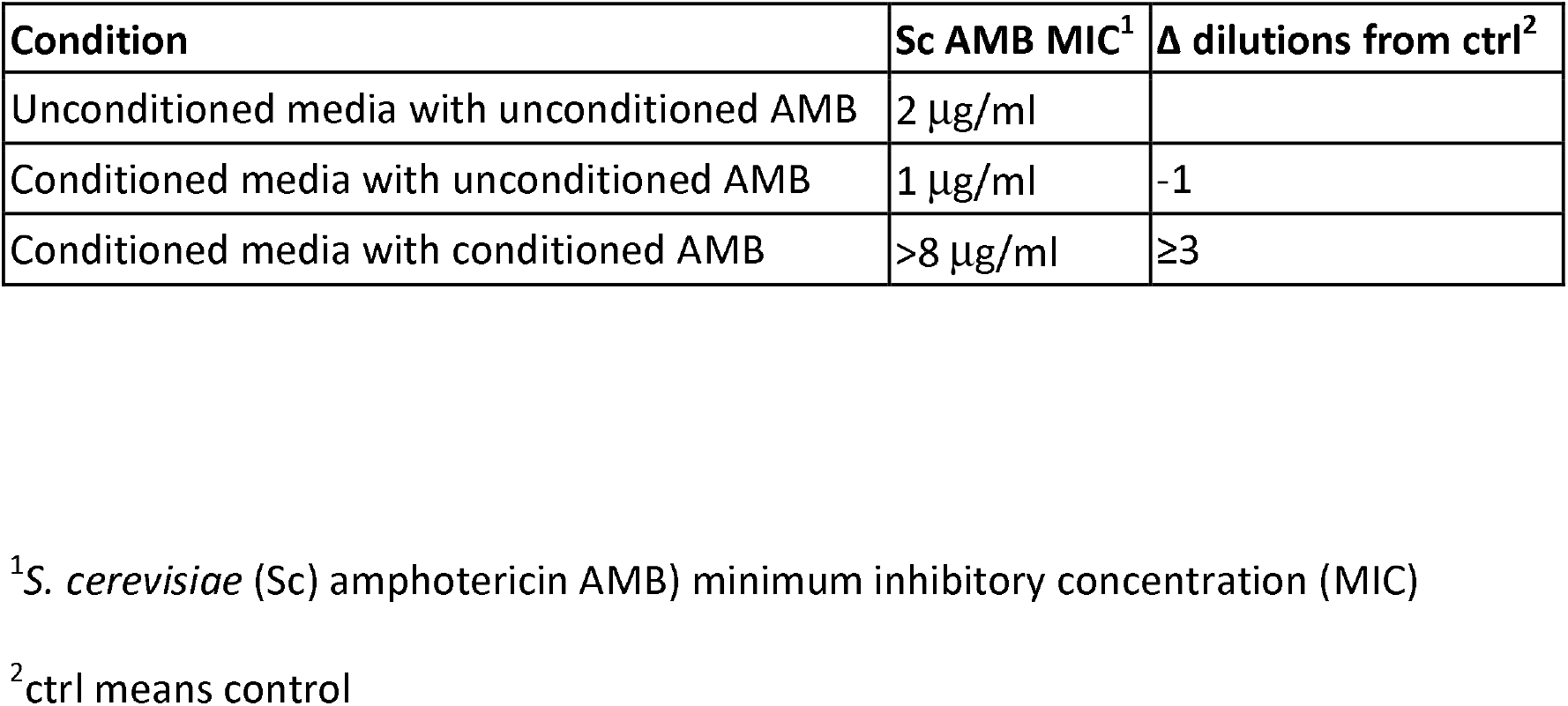
Drug activity assay results of RPMI incubated with *L. prolificans* before and after addition of AMB.

Given that the initial discrepancy in AMB susceptibility between *L. prolificans* conidia and protoplasts suggested a role for the cell wall in drug resistance, we tested whether the dramatic difference between the abilities of *L. prolificans* and *A. fumigatus* to alter drug activity would be maintained as protoplasts. By wet pellet mass, 4 mg/ml of *L. prolificans* protoplasts in media diminished drug activity by one dilution, while 8 mg/ml of *A. fumigatus* was necessary to produce the same effect (Table 3). By comparison, intact *L. prolificans* was about eight times as potent as intact *A. fumigatus* at reducing drug activity. Interestingly, the two-fold difference in the drug activity-reducing capacity of *L. prolificans* and *A. fumigatus* protoplasts mirrors the single dilution difference in AMB susceptibility between *L. prolificans* and *A. fumigatus* protoplasts. The reduction of the effect of *L. prolificans* on drug activity compared to that of *A. fumigatus* following the removal of the cell wall reinforces the suggestion that the source of *L. prolificans* AMB resistance resides in the cell wall.

**Table 3.** Drug activity assay results of AMB incubated with *L. prolificans* (Lp) or *A. fumigatus* (Af) protoplasts.

| Species/conc. |  | Sc AMB MIC | Δ dilutions from ctrl<sup>2</sup> |
| --- | --- | --- | --- |
| No treatment |  | 1 µg/ml |  |
| Lp protoplasts | 2 mg/ml | 1 µg/ml | 0 |
|  | 4 mg/ml | 2 µg/ml | 1 |
|  | 8 mg/ml | 4 µg/ml | 2 |
| Af protoplast | 4 mg/ml | 1 µg/ml | 0 |
|  | 8 mg/ml | 2 µg/ml | 1 |

<sup>1</sup>*S. cerevisiae* (Sc) amphotericin (AMB) minimum inhibitory concentration (MIC)
<sup>2</sup>ctrl means control

In C. albicans biofilms, β-glucans have been linked to resistance to both AMB and fluconazole (9, 10, 12). To determine whether the same phenomenon could be occurring in *L. prolificans*, we tested whether incubation with *L. prolificans* could diminish fluconazole activity in the same manner it does AMB activity. Incubation with *L. prolificans* had no effect on fluconazole activity (data not shown).

Having established that *L. prolificans* can absorb or otherwise inactivate AMB, and that the cell wall is essential to this capacity, we evaluated whether cells must be alive and intact to produce this effect, or if the mere presence of the cellular materials was sufficient. Accordingly, the drug activity assay was conducted in parallel using an equal mass of either homogenized or intact cells. The resulting drug microdilution plate showed the same amount of growth in the homogenate- and intact cell-incubated media, demonstrating that the two diminished AMB activity to the same extent (Table 4). This demonstrated that the effect on drug activity was not contingent on live, or even intact, *L. prolificans*.

**Table 4.** Drug activity assay results of AMB incubated with intact or homogenized *L. prolificans* (Lp) hyphae.

| Species/concentration | Sc AMB MIC<sup>1</sup> | $\Delta$ dilutions from ctrl<sup>2</sup> |
| --- | --- | --- |
| No treatment | 2 μg/ml | |
| Lp intact cells (1 mg/ml) | >8 μg/ml | $\geq 3$ |
| Lp homogenized cells (1 mg/ml) | >8 μg/ml | $\geq 3$ |

<sup>1</sup>*S. cerevisiae* (Sc) amphotericin AMB) minimum inhibitory concentration (MIC)
<sup>2</sup>ctrl means control

To pursue the suggestion that a cellular component, as opposed to a cellular process, was responsible for abrogating the AMB drug activity, the *L. prolificans* cell homogenate was centrifuged and separated into three fractions: supernatant, a light-colored top layer of pellet and a darker bottom layer of pellet. Each of these was then used in place of hyphae in the drug activity assay, with concentration normalized by the sample’s starting hyphal mass. The supernatant had the same drug-diminishing effect as the unfractionated homogenate, while the top layer of pellet showed no effect on drug activity, and the bottom layer of pellet showed only minor effects on drug activity (Table 5). For comparison, the same experiment was conducted with an equal mass of *A. fumigatus* and none of the fractions showed any effect on drug activity. This was somewhat surprising, given the previous findings demonstrating the involvement of the cell wall in diminishing AMB activity, as most components of the cell wall would be expected to be present in the pellet, rather than the supernatant. The finding suggests a soluble component attached to or contained within the cell wall but released by homogenization is responsible for the observed reduction in AMB activity.

**Table 5.** Drug activity assay results of AMB incubated with *L. prolificans* (Lp) or *A. fumigatus* (Af) homogenate fractions.

| Species/fraction (1 mg starting culture/ml) | Sc AMB MIC<sup>1</sup> | Δ dilutions from ctrl<sup>2</sup> |
| --- | --- | --- |
| No treatment | 1 µg/ml |  |
| Lp homogenate supernatant | >8 µg/ml | ≥4 |
| Lp homogenate pellet – top layer | 1 µg/ml | 0 |
| Lp homogenate pellet – bottom layer | 2 µg/ml | 1 |
| Af homogenate supernatant | 1 µg/ml | 0 |
| Af homogenate pellet – top layer | 1 µg/ml | 0 |
| Af homogenate pellet – bottom layer | 1 µg/ml | 0 |

<sup>1</sup>*S. cerevisiae* (Sc) amphotericin (AMB) minimum inhibitory concentration (MIC)
<sup>2</sup>ctrl means control

To further elucidate the attributes of this “resistance factor,” the homogenate supernatant was fractionated by size. Passing through a 0.22 μm filter prior to being used in the drug activity assay had no effect on efficacy compared to supernatant that was not prefiltered. Following this, all future drug activities were performed such that the cellular component of interest was added to media, then filter-sterilized, and then aliquoted for the addition of drug. This eliminated the need for centrifugation and filtration of individual samples (Step 4 in Figure 1).

Further fractionation was performed using Amicon filters ranging from 3 kda to 100 kda in size, and the resulting fractions were used in a drug activity assay. Of these, only the >100 kda fraction maintained an inactivating effect on AMB (Table 6).

**Table 6.**
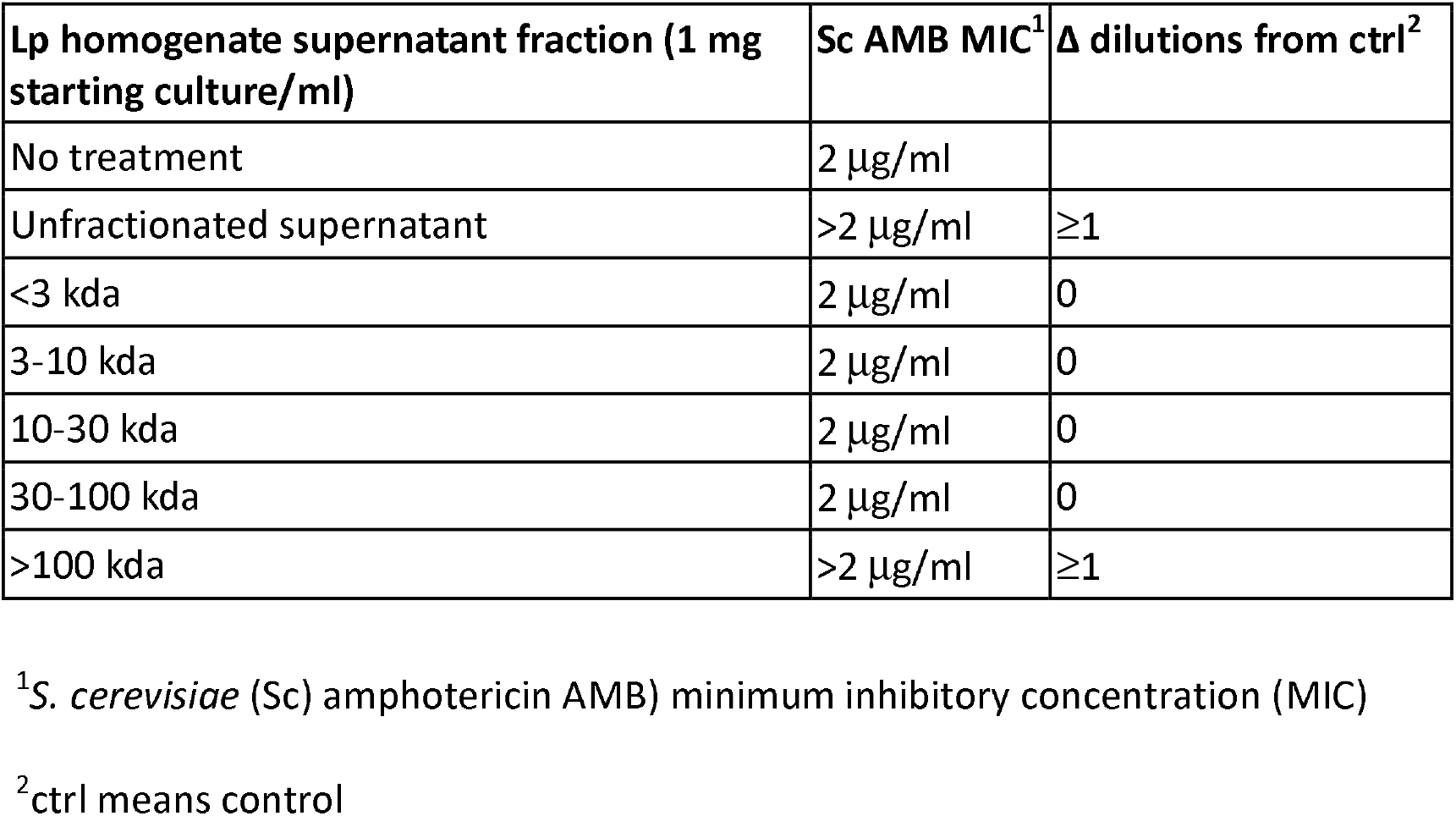
Drug activity assay results of AMB incubated with *L. prolificans* homogenate supernatant fractions fractionated by size filtration.

To further test the hypothesis that the resistance factor was contained within the cell wall, as suggested by the previous protoplast experiments, we tested the effects on drug activity of the supernatant and pellet left over after the generation of protoplasts by treatment with cell wall-digesting enzymes. While the pellet, which was divided into three layers by color, each of which was tested separately, showed no effect of AMB activity, the supernatant impaired drug activity to the same extent as whole cell homogenate supernatant, normalized by starting culture weight (Table 7). Together, this bolsters the suggestion that the resistance factor resides within the cell wall and is soluble in the supernatant upon being released either physically (through homogenization) or chemically (through cell wall digestion).

**Table 7.** Drug activity assay results of AMB incubated with fractions of material remaining the generation of *L. prolificans* protoplasts.

| **Treatment** | **Sc AMB MIC<sup>1</sup>** | **Δ dilutions from ctrl<sup>2</sup>** |
| --- | --- | --- |
| No treatment | 2 µg/ml |  |
| Unfractionated supernatant | >2 µg/ml | ≥1 |
| Post-protoplasting supernatant | >2 µg/ml | ≥1 |
| Post-protoplasting pellet – top layer | 1 µg/ml | -1 |
| Post-protoplasting pellet – middle layer | 1 µg/ml | -1 |
| Post-protoplasting pellet – bottom layer | 1 µg/ml | -1 |

<sup>1</sup>*S. cerevisiae* (Sc) amphotericin AMB) minimum inhibitory concentration (MIC)
<sup>2</sup>ctrl means control

A series of treatments were tested for their ability to eliminate the action of the resistance factor. Treatment with DNase, trypsin, proteinase k, amyloglucosidase, zymolyase and SDS had no impact on the action of the resistance factor (Table 8). However, treatment with urea or acetonitrile prior to incubation in media for the drug activity assay eliminated the resistance factor’s effect, as did heating to 65°C for one hour. Additionally, attempts to separate the sample into lipid and non-lipid fractions using two different methods resulted in no resistance factor effect in any resulting fraction.

**Table 8.** Inactivation treatments grouped by their effect on the capacity of Lp supernatant to counteract AMB activity.

**AMB activity**
| **No difference**<br>**(MIC<sub>treated supernatant</sub> = MIC<sub>untreated supernatant</sub>)** | **Complete elimination**<br>**(MIC<sub>treated supernatant</sub> = MIC<sub>no supernatant</sub>)** |
| --- | --- |
| DNase | Urea |
| Trypsin | BUME organic phase |
| Proteinase K | BUME aqueous phase |
| Amyloglucosidase | Acetonitrile (60%) |
| SDS | Methanol/chloroform organic phase |
| 0.22 µm filtration | Methanol/chloroform aqueous phase |
|  | Heat (65°C for 1 hr) |

## Discussion

In this paper, we report on the existence of a cell wall substance in *L. prolificans* that can eliminate the activity of AMB. This represents the first mechanism of AMB resistance identified in *L. prolificans*. While the mechanism by which the resistance substance interferes with AMB is unclear, it likely binds to the drug and prevents its binding to ergosterol.

While not definitive, our findings are consistent with the substance being a glycolipid. The suggestion is supported by the fact that, while most means of inactivation by chemical or enzymatic treatment were unsuccessful, separation into aqueous and organic phases eliminated activity entirely. Previous work has shown that β-glucans are capable of facilitating AMB resistance, apparently by absorbing the drug and preventing it from reaching the cell membrane. It appears less likely that β-glucans are involved in this case, given that treatment with multiple β-glucanase enzymes did not diminish the drug resistance factor’s effects on AMB, and indeed digesting *L. prolificans* with lysing enzymes from Trichoderma harizianum served to release the factor into solution in an equivalent manner to homogenization. However, it is possible that the nature of the interaction between AMB and β-glucan, a polysaccharide, is similar to the interaction between AMB and the polysaccharide component of the resistance factor.

The revelation that the presence of *L. prolificans* is sufficient to reduce AMB activity has implications for clinical treatment. While *L. prolificans* rarely causes invasive infections, its presence in the lung appears to be more common, particularly in cystic fibrosis patients (25, 26). It is plausible that, even in a commensal state, *L. prolificans* may actively protect other pathogenic fungi from clearance with AMB treatment.

It is noteworthy that *S. apiospermum* displayed an ability to inactivate AMB in media that was considerably less than that of *L. prolificans*, but considerably greater than that of *A. fumigatus*, as, indeed, its susceptibility to AMB is between that of the two other species. It is possible that Scedosporium species trace their unusual AMB tolerance to the generation of a similar substance, either to a lesser extent than *L. prolificans* or in a less effective form.

In summary, AMB resistance in *L. prolificans* appears to be mediated by a soluble cell wall resistance factor. The factor appears to be a macromolecule with a mass > 100 Kd but we don’t know if it is a single molecule or a complex. We note that this mechanism of AMB resistance is different from others described, implying the existence of cell wall substances that protect against this drug. While we were unable to purify this substance to a purity sufficient for structural studies, we note that its large size and likely heterogeneity could pose a formidable analytical problem. Nevertheless, the work described here provides a roadmap for future analytical studies and sets a precedent for the existence of new mechanisms of AMB resistance.

## Materials and Methods

### Strains

The strains used in this work were *L. prolificans* strain 3.1 from Christopher Thornton, *Scedosporium apiospermum* strain IHEM 14462 from BCCM, Aspergillus fumigatus strain MYA-3626 and *Saccharomyces cerevisiae* strain S288C from the Susan Michaelis lab. Conidia were obtained from the filamentous fungi by inoculating frozen conidia into sabouraud dextrose broth, growing this up for 5-21 days shaking at 30°C, then plating 2-3 ml on potato dextrose agar. After growing for 6-8 days, these plates were covered in DPBS without magnesium or calcium and scraped. The resulting conidia in DPBS were strained through a 70 μm cell strainer, then pelleted at 4000 rpm and washed three times. In the case of *A. fumigatus*, 0.05% tween 20 was added to DPBS, which was then passed through a 0.22 μm Steritop filter (Millipore Sigma, Burlington, MA).

Cultures were grown by inoculating conidia into RPMI to a concentration of 10^6^ conidia/ml. These were grown shaking at 90 RPM for 16 hours at 37°C, then strained through two layers of miracloth (Millipore Sigma) and washed with Milli-Q water. Fungal biomass was squeezed dry with Kimwipes (Kimberly-Clark, Irving, TX) and weighed.

### Protoplasts

Protoplasts were generated using published methodology with minor modifications (21). Briefly, conidia were inoculated into sabouraud dextrose broth to a concentration of 10^6^ conidia/ml and shaken at 30°C for ~16 hours. This culture was strained through two layers of sterile miracloth, washed with sterile milli-Q water, and then the fungal biomass was transferred to sterile-filtered OM buffer with lysing enzymes from *Trichoderma harizianum* (1.2 M MgSO4 - 7H2O, 10 mM NaPO4 (pH 5.8), 5% lysing enzymes by weight) and shaken at 30°C overnight. Digested cells in OM buffer were transferred into Oak Ridge centrifuge tubes, overlaid with chilled ST buffer (0.6 M Sorbitol, 10 mM Tris-HCl (pH 7.0)), and centrifuged at 5000 x g 4°C for 15 minutes in a swinging bucket rotor. Protoplasts were recovered at the OM/ST interface, transferred to an Oak Ridge centrifuge tube with cold STC buffer (1.2 M sorbitol, 10 mM Tris-HCl (pH 7.5), 10 mM CaCl_2_), and pelleted at 3000 x g for 10 minutes 4°C. Protoplasts were then washed and pelleted twice more.

### Antifungal susceptibility testing

Antifungal susceptibility testing was carried out using CLSI broth microdilution methodology (27). Briefly, conidia or protoplasts were collected as described above and diluted into RPMI with 1.2 M sorbitol and inoculated 1:1 into broth microdilution plates to a final concentration of 2×10^5^ (protoplasts) or 5 x 10^3^ (conidia) cells/ml and 0.6 M sorbitol. Plates were grown at 37°C in 5% CO_2_ for three days, then imaged. To ensure that the growth in tests of protoplast susceptibility reflected only protoplasts, and not any remaining intact cells, protoplasts were also diluted into RPMI without sorbitol and then inoculated into broth microdilution plates.

### Drug activity assay

The extent to which drug activity was reduced by the presence of fungal biomass or fractions thereof was assessed as follows. A known concentration of the substance of interest was added to RPMI, which was then divided into aliquots to which a 100x dilution of AMB in DMSO was added to a final 1x concentration of AMB. These samples were then shaken at 37°C for one hour. In experiments in which the substance of interest could pass through a 0.22 μm filter, RPMI with the substance was sterile filtered prior to aliquoting and adding drug. In experiments in which the substance could not pass through a filter, such as when using intact fungal cells or fungal homogenate, samples were sterile filtered individually after shaking. S. cerevisiae strain S288C was then added to each sample to a final concentration of 1.5×10^3^ cells/ml and samples were transferred to a U-bottom 96-well plate, which was incubated at 30°C for 2-3 days.

### Cell homogenization

Fungal biomass that had been squeezed dry and weighed was resuspended in Milli-Q water and added to screw-top tubes with 0.5 mm glass beads. This was shaken in a FastPrep-24 bead beater (MP Biomedicals, Irvine, CA) five times at 6 m/s for one min, with 1.5 min on ice in between. Fractionation of the supernatant was conducted by microcentrifugation for 15 minutes at 4°C.

### Digestions, treatments and fractionation of homogenate supernatant

DNase digestion was conducted using PureLink DNase I (Invitrogen, Waltham, MA) reconstituted per manufacturer’s instructions. Reaction consisted of 80 μl of supernatant, 10 μl of DNase buffer (as provided), and 10 μl of DNase I. This was incubated at room temperature for 15 min. Trypsin digestion was performed using 7.5 μl of 0.05% Trypsin-EDTA (ThermoFisher Scientific, Waltham, MA) with 100 μl of supernatant. Reaction was shaken at 37°C for 30 min, then halted using a final concentration of 0.033% soy bean trypsin inhibitor (Millipore Sigma). SDS treatment was conducted using a final concentration of 0.1% SDS. SDS was then removed from sample using a 100 kda Amicon filter (Millipore Sigma), and retained fraction was used in drug activity assay. Urea treatment was conducted at a final concentration of 4 M urea. Amyloglucosidase digestion was conducted using 0.2% by volume aqueous solution of amyloglucosidase from Aspergillus niger in 25 mM sodium acetate at 37°C for 30 min. Proteinase K digestions were conducted with 0.2 mg/ml recombinant, PCR-grade proteinase K (Roche, Basel, Switzerland) in 30 mM Tris-HCl (pH 8.3) at shaking at 37°C for 1 hr. Where specified, digestions were carried out instead with 10 mg/ml proteinase K-agarose. Digestion with zymolyase 20T (ThermoFisher Scientific) was performed at 0.05 mg/ml of enzyme in M/15 buffer for 1 hr at 37°C.

### Lipid extraction

Two different lipid extraction methods were utilized. The first was adapted from the Bligh and Dyer method (28). Briefly, 750 μl of 1:2 chloroform:methanol was added to 200 μl of sample and vortexed for 10 min. Then 250 μl of chloroform was added, and the sample was vortexed for 1 min. Finally, 250 ul water was added, sample was vortexed for 1 min and phases were collected. The second approach was the BUME method developed by Löfgren et al. in 2012 (29). The procedure was performed as written, with the only alteration that it was carried out by hand, rather than by robot.

## Acknowledgments

NTG was supported in part by T32AI007417. A.C. was supported in part by National Institutes of Health R01 Grants HL059842, AI152078, AI052733 and U19AI189183.

